# A silicon rhodamine-fused glibenclamide to label and detect malaria-infected red blood cells

**DOI:** 10.1101/2025.02.07.637098

**Authors:** Claudia Bastl, Cindy M. Close, Ingo Holtz, Blaise Gatin-Fraudet, Mareike Eis, Michelle Werum, Smilla Konrad, Kilian Roßmann, Christiane Huhn, Souvik Ghosh, Julia Ast, Dorien A. Roosen, Martin Lehmann, Volker Haucke, Luc Reymond, David J. Hodson, Philip Tinnefeld, Kai Johnsson, Viktorija Glembockyte, Nicole Kilian, Johannes Broichhagen

**Affiliations:** Centre for Infectious Diseases, Parasitology Heidelberg University Hospital, Im Neuenheimer Feld 324, 69120 Heidelberg, Germany; Department of Chemistry and Center for NanoScience (CeNS), Ludwig-Maximilians-University, Butenandtstraße 5–13, 81377 Munich, Germany; Leibniz-Forschungsinstitut für Molekulare Pharmakologie, Robert-Rössle-Straße 10, 13125 Berlin, Germany; Institute of Metabolism and Systems Research (IMSR), and Centre of Membrane Proteins and Receptors (COMPARE), University of Birmingham, Birmingham, UK; Department of Molecular Pharmacology and Cell Biology, Leibniz-Forschungsinstitut für Molekulare Pharmakologie, Robert-Rössle-Straße 10, 13125 Berlin, Germany; Laboratory of Protein Engineering, Institut des Sciences et Ingénierie Chimiques, Sciences de Base, École Polytechnique Fédérale Lausanne, Avenue Forel 2, 1015 Lausanne, Switzerland; Biomolecular Screening Facility, École Polytechnique Fédérale Lausanne, Avenue Forel 2, 1015 Lausanne, Switzerland; Oxford Centre for Diabetes, Endocrinology and Metabolism (OCDEM), NIHR Oxford Biomedical Research Centre, Churchill Hospital, Radcliffe Department of Medicine, University of Oxford, Oxford, UK; Department of Chemical Biology, Max Planck Institute for Medical Research, Jahnstr. 29, 69120 Heidelberg, Germany; Department of Biological Sciences, California State University, Chico, USA, 400 W. First St. Chico, CA 95929–515, USA; Department of Medical Biochemistry, Faculty of Basic Medical Sciences, Delta State University, P.M.B. 1 Abraka Delta State, Nigeria

**Keywords:** Silicon Rhodamine, Red Blood Cells, Malaria, *Plasmodium falciparum*, Smartphone-Based Detection

## Abstract

The malaria parasite *Plasmodium falciparum* affects the lives of millions of people worldwide every year. The detection of replicating parasites within human red blood cells is of paramount importance, requiring appropriate diagnostic tools. Herein, we design and apply a silicon rhodamine-fused glibenclamide (**SiR-glib**). We first test this far-red fluorescent, fluorogenic and endoplasmic reticulum-targeting sulfonylurea in mammalian cells and pancreatic tissues, before characterizing its labeling performance in red blood cells infected with the asexual developmental stages of *Plasmodium falciparum*. We further combine **SiR-glib** with a portable smartphone-based microscope to easily and rapidly identify parasitized red blood cells, providing proof of principle for diagnostic use in rural endemic areas without major healthcare facilities.

## Introduction

The latest World Malaria Report published by the World Health Organization (WHO) in December 2022 recorded 247 million malaria associated clinical cases and 619,000 deaths^[1]^. Malaria tropica, the major and most dangerous form of human malaria, is caused by the protozoan parasite *Plasmodium falciparum (P. falciparum*) and is responsible for maternal illness, low birth weight and patient deaths in endemic areas^[1–3]^.

Detection of *P. falciparum* is important to initiate the appropriate anti-malarial therapy, especially in the field where access to healthcare facilities, electricity and transportation might be restricted^[1,4]^. We therefore started with the premise that mature human red blood cells (RBCs), which serve as a host cell during the parasite’s asexual reproduction (erythrocytic schizogony), do not contain any organellar structures (i.e. they are anuclear) (**Figure 1A**). *P. falciparum*, on the other hand, contains cell organelles that i) are found in other eukaryotic cells (e.g. endoplasmic reticulum, Golgi apparatus, nucleus); and ii) are specialized, such as the apicoplast and the digestive vacuole. The parasite interacts and reorganizes the cell organelle-deprived host RBCs to acquire nutrients, ultimately maturing from a young ring to a trophozoite and finally to a schizont stage within approximately 48 hours^[5–7]^.

**Figure 1.**
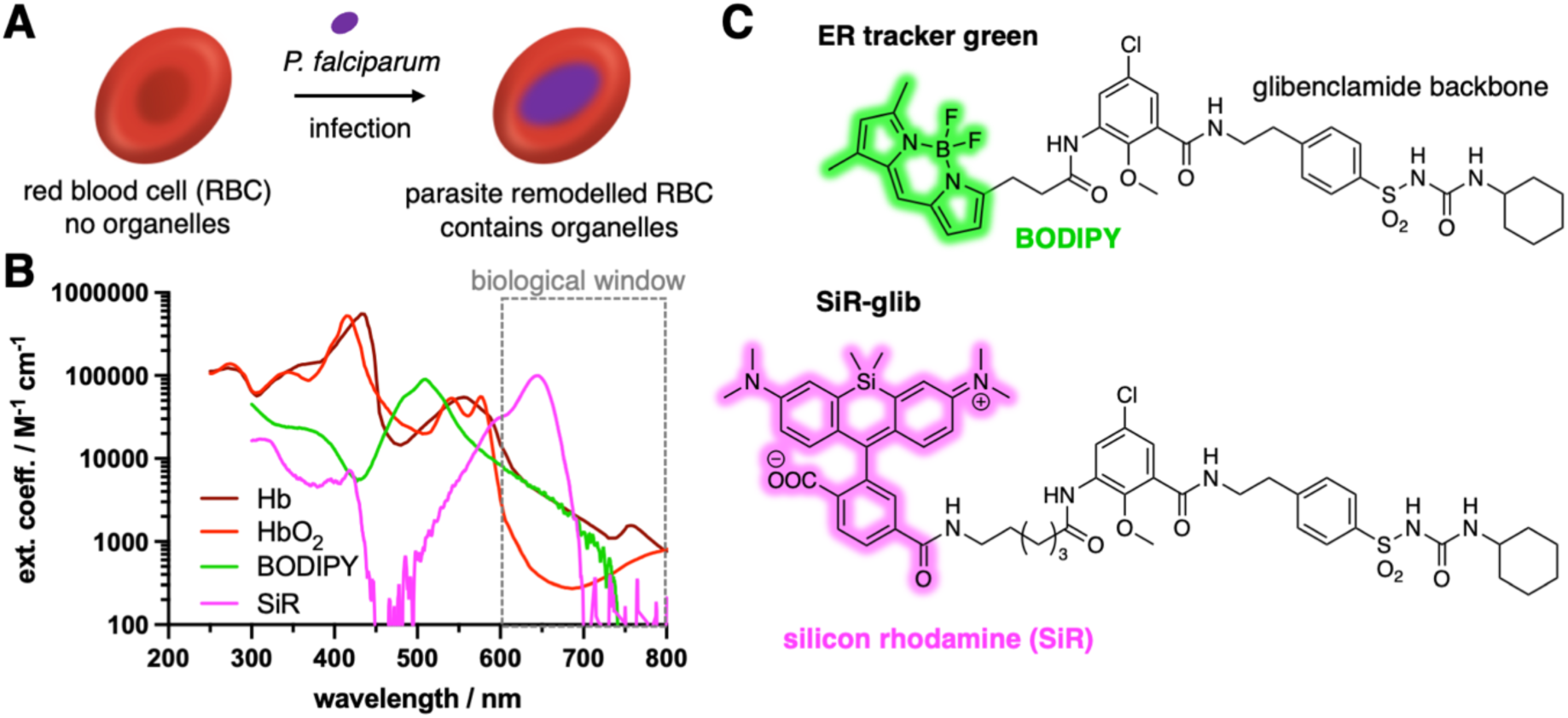
Silicon rhodamine-fused glibenclamide for the detection of *P. falciparum* infected red blood cells (RBCs). **A)** RBC do not contain any organelles, which changes through the infection by *P. falciparum* parasites due to remodeling and genesis of ER, nucleus, etc. **B**) High absorbance of oxygen free (Hb) and oxygen-bound hemoglobin (HbO_2_) in the UV to green range might mask ER tracker green. Far-red silicon rhodamine falls into the biological imaging window above 600 nm. **C**) Chemical structure of glibenclamide and its fluorophore-linked congeners ER tracker green (BODIPY fused) and **SiR-glib** (SiR fused).

We focused our investigation on the cell organelles of the parasite that may be visualized via fluorescence microscopy with small molecule chemical biology probes^[8]^. Potential candidates such as ER trackers Green/Red (λ_ex_ = 504 nm or 588 nm), which comprise glibenclamide-BODIPY targeting the sulfonylurea receptor 1, are of limited use as they fall into the same spectral window as (non)-oxygenated hemoglobin (Hb and HbO_2_) (**Figure 1B**). Far-red and near-infrared fluorophores are well-suited to use in RBC’s, since blood displays an optical window between 600–800 nm where absorbance and hence background fluorescence is low.^[9]^ We therefore decided to design an ER tracker using silicon rhodamine (SiR) as a fluorogenic far-red dye, chemically fused it with glibenclamide and named it **SiR-glib**. The probe was applied across different imaging contexts to stain the ER in live mammalian cells and tissue, and in infected RBCs (iRBCs). To detect infected stages, we used a home-built, cheap smartphone-based microscope^[10]^, which bears the potential for malaria diagnostic applications in endemic areas that have poor access to healthcare facilities and electricity.

## Results

We set out to conjugate far-red silicon rhodamine (SiR)^[11]^, a dye previously used in RBCs for actin staining^[12]^, to the sulfonylurea glibenclamide^[13]^, which targets SUR1 expressed on the endoplasmic reticulum (ER). The synthesis starts by activating 5-chloro-2-methoxy-3-nitrobenzoic acid (**1**) using TSTU and forming a peptide bond with 4-(2-aminoethyl)benzenesulfonamide to obtain sulfonamide **2** in 81% yield (**Scheme 1**). Using cyclohexyl isocyanate in acetone with K_2_CO_3_ serving as a base, the sulfonylurea motif of **3** was installed in quantitative yield. Reduction of the nitro group using zinc and acetic acid in methanol progressed in 90% yield, and the corresponding aniline **4** was endowed with a 6-carbon atom long linker. To address the rather unreactive aniline, we first formed an acyl chloride *in situ* by stirring Fmoc-Ahx-OH in neat SOCl_2_. After evaporation of all volatiles and reuptake in DIPEA containing 1,4-dioxanes, the acyl chloride was added to a solution of **4**. The solvents were evaporated, and the crude material containing **5** taken up in DMF with 5% piperidine to deprotect the Fmoc group, allowing the isolation of alkyl amine **6** in 17% yield over this synthetic sequence. Finally, we obtained a fluorogenic (**Figure 2A**) silicon rhodamine-fused glibenclamide, termed **SiR-glib**, in 42% yield after HPLC purification by using NHS-activated ester of silicon rhodamine. We first confirmed the expected spectral properties (λ_ex_/λ_em_ = 653/673 nm) (**Figure 2B**) of **SiR-glib**, and tested its fluorogenicity and pH-sensitivity. **SiR-glib** displayed a 10.0-fold fluorescence increase when SDS was added to the buffer medium (**Figure 2B**), and a trend to decreased emission at more acidic pH values (**Figure 2C**). This is warranted because iRBCs maintain a physiological pH while the digestive vacuole, which is responsible for hemoglobin digestion and storage of hemozoin has been shown to exhibit a drop in pH to ∼ 5.2 (ref^[14,15]^).

**Figure 2.**
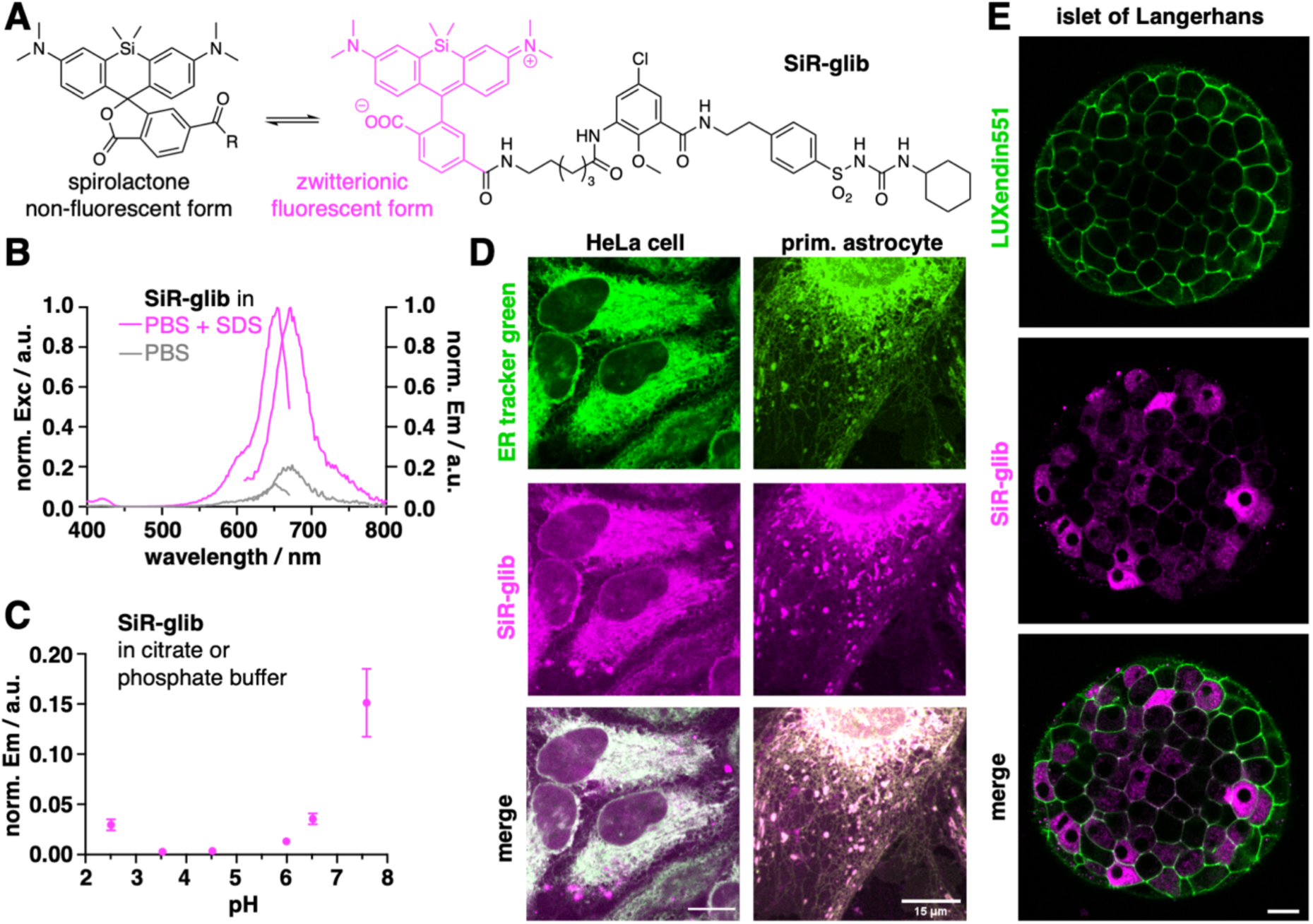
Characterization of SiR-glib. **A) SiR-glib** may adopt two different isomeric forms, a non-fluorescent spirolactone and an open, fluorescent zwitterionic form. **B)** Excitation and emission spectra of **SiR-glib**, showcasing its fluorogenicity by addition of SDS. **C**) Fluorescence pH dependency of **SiR-glib**. **D**) Live cell imaging by confocal microscopy using ER tracker green and **SiR-glib** in live HeLa cells and live primary astrocytes. **E**) Live cell imaging by confocal microscopy beta-cell marker LUXendin551 and **SiR-glib** in live islets of Langerhans. **SiR-glib** gives rise to stained membrane structures, presumably due to KATP binding. Scale bars = 15 µm.

**Scheme 1.**
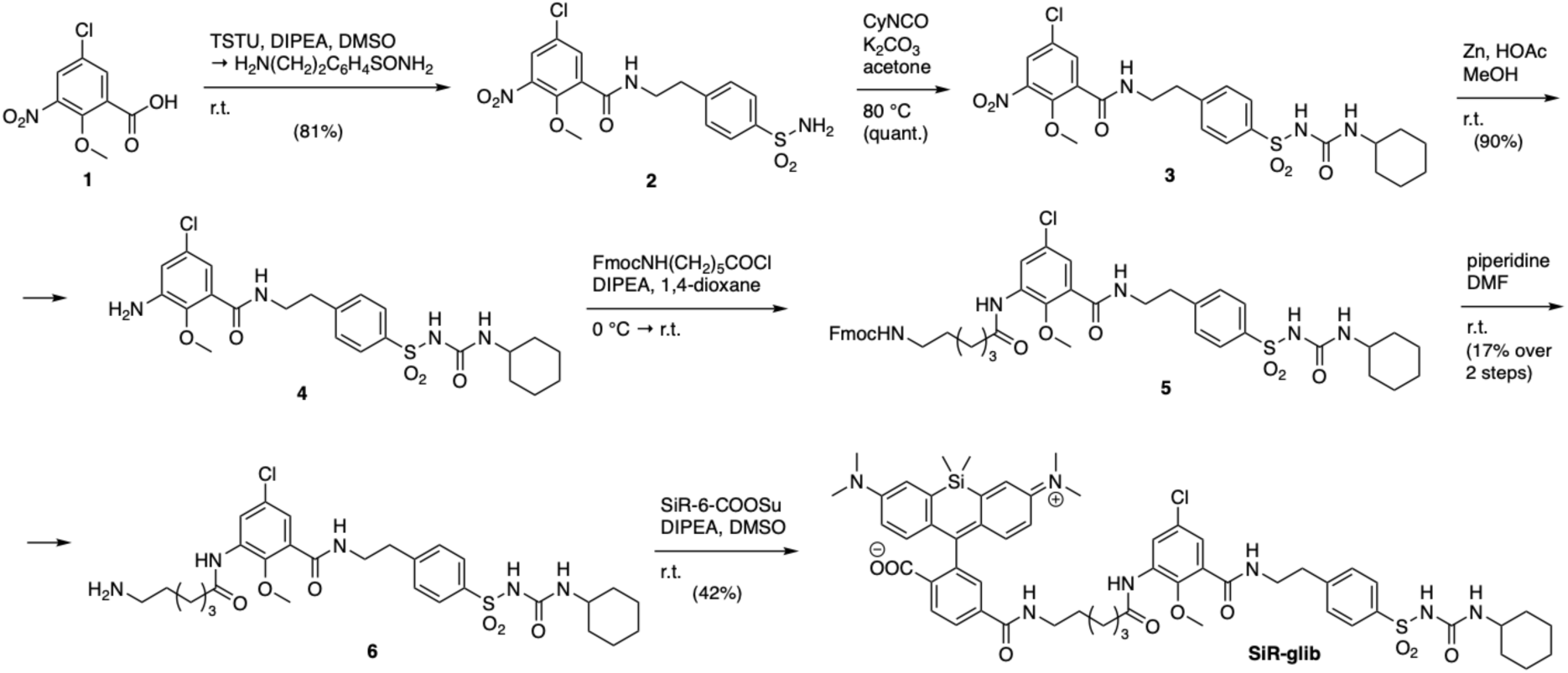
Synthesis of SiR-glib. Commercially available benzoic acid **1** is peptide coupled after TSTU activation to obtain **2**, on which the sulfonylurea is installed using cyclohexyl isocyanate. Zn-mediated reduction of the nitro group yields aniline **4**, on which an alkyl amine linker is installed using an *in situ* formed acyl chloride before subsequent Fmoc-deprotection. Silicon rhodamine is finally fused onto **6** by using its NHS-activated ester, yielding **SiR-glib**.

Next, we set out to determine the performance of **SiR-glib** in live cell imaging by incubating HeLa cells with 5 µM ER tracker green and 50 µM **SiR-glib**, before imaging by confocal fluorescence microscopy (**Figure 2D**). By merging the images, we found good co-localization between ER tracker green and **SiR-glib**, showcasing accurate targeting of the ER by **SiR-glib**. While immortalized cell lines can also be marked with fluorescent proteins containing appropriate targeting sequences (e.g., Sec61^[16]^), genetic engineering is more difficult in primary cells, let alone in parasites or even patient samples. Accordingly, we compared ER tracker green and **SiR-glib** in primary mouse astrocytes, and were able to acquire similar staining patterns.

Since glibenclamide targets the sulfonylurea receptor 1 (SUR1), which is retained at the ER in the absence of Kir6.2, we decided to test **SiR-glib** in cells that express both SUR1 and Kir6.2. We reasoned that in cells with both subunits-i.e. those that express K_ATP_ channels-staining should be seen within the cell (ER, SUR1) and at the membrane (SUR1 + Kir6.2 octamer i.e. KATP channel).^[17]^ Islets of Langerhans were thus incubated with 100 nM of the fluorescent beta cell marker LUXendin551^[18]^ and 50 µM **SiR-glib**, before confocal live imaging. **SiR-glib** staining was seen both within the cell (SUR1) as well as at the cell membrane (SUR1 + Kir6.2), further demonstrating specificity of the probe.

Encouraged by this, we applied **SiR-glib** to *in vitro* cultures of *P. falciparum*-iRBCs. We chose two standard laboratory model strains for our investigations, 3D7 (clone of NF54, originating from an airport malaria case) and FCR3 (originating from Gambia)^[19–22]^. *In vitro* cultures were infected with *P. falciparum* and incubated with either 2 µM **SiR-glib**, 2 µM SiR or remained untreated for 1 h at 37 °C at 5% hematocrit (HCT, volume percentage of RBCs in the human body or in this case, in the in vitro culture). Investigation by confocal laser scanning microscopy revealed distinct labeling of all developmental stages (i.e., ring, trophozoite and schizont) of 3D7 (**Figure 3A**) and FCR3 (**Figure 3B**) *P. falciparum* laboratory strains. More specifically, a punctuate signal was observed in ring stages, whereas in trophozoite and schizont stages a line or circle near or around the digestive vacuole, respectively, was eminent (**Figure 3A, B**). We quantified the fluorescence signal of iRBCs and observed significant differences between **SiR-glib**-, SiR- or non-treated controls of 3D7 (**Figure 3C**) and FCR3 (**Figure 3D**). These results show that the glibenclamide scaffold is needed for successful organellar targeting and yields an 11.5-fold (3D7) and 8.0-fold (FCR3) increase in fluorescence signal in all stages of iRBCs. To test if the parasites may be properly labeled in a higher hematocrit, we prepared a 40% HCT culture and inoculated it with the *P. falciparum* lab strains 3D7 and FCR3. This is particularly important because HCT of human blood for females and males ranges from 36– 48% and 40–54%, respectively^[23]^. After allowing the parasites to adapt and replicate for 48 h, the culture was incubated with **SiR-glib** and SiR as before, or kept untreated for 1 h at 37 °C. Again, successful labeling of the different developmental stages with **SiR-glib** was observed, which differed from the control treatments of the two *P. falciparum* cultures (**Figure 3E, F**). While a loss in fluorescence signal was observed in samples containing ring stages, the overall fluorescence increases for all stages in the infected samples remained significantly high (9.6-fold for 3D7 and 4.2-fold for FCR3). These experiments highlight the sensitivity of **SiR-glib** in dilute samples, as well as in whole blood model systems.

**Figure 3.**
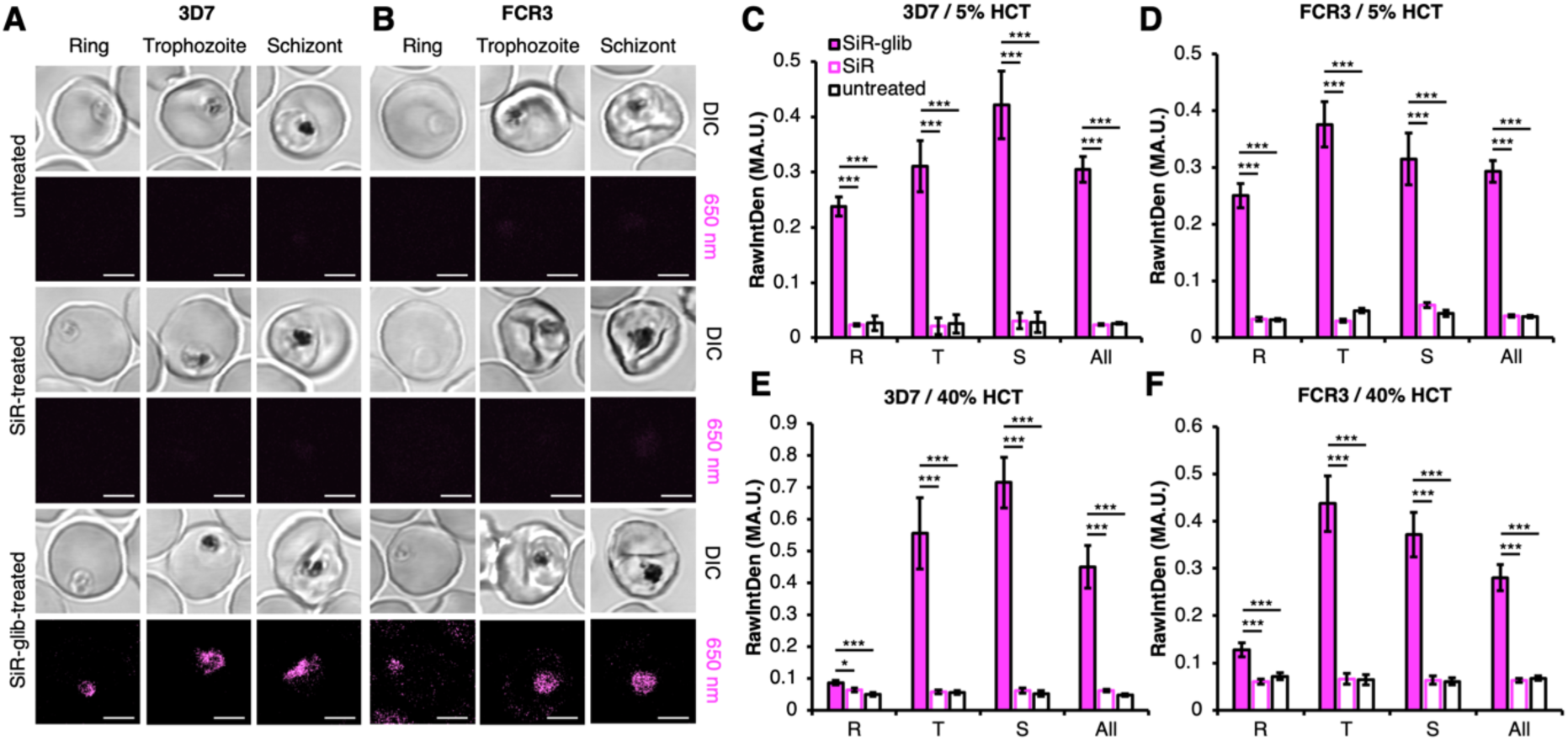
Labeling of two different *P. falciparum* strains with SiR-glib. **A)** The *P. falciparum* strain 3D7 was grown in a 5% HCT culture. Cells were labeled with 2 µM **SiR-glib**, 2 µM SiR or remained untreated for 1 h. Only **SiR-glib**-treated cells showed a specific labeling of the ring (R), trophozoite (T) and schizont (S) developmental stages of the parasite for both investigated strains. Signals of all stages are included (All). Scale bars = 3 µm. **B)** As for (**A**) but using the *P. falciparum* strain FCR3. **C, D)** Quantification of fluorescence from (A) and (B) reveals a significantly stronger fluorescence of **SiR-glib**-treated RBCs compared to using SiR or untreated RBCs in every developmental stage. N=3; n≥11. *: p < 0.05; **: p < 0.01; ***: p < 0.001. **E, F)** As for (**C**) and (**D**) but in a 40% HCT culture shows lower signal in ring stages when compared to 5% HCT. N=3; n≥12. *: p < 0.05; **: p < 0.01; ***: p < 0.001.

Finally, we were wondering if **SiR-glib** could potentially be used to label *P. falciparum* in point-of-care diagnostic settings in endemic areas^[4]^. Therefore, the treatment of the *in vitro* cultures was adjusted: the concentration of **SiR-glib** was increased from 2 µM to 50 µM, the incubation time was shortened from 1 h to 10 min and the incubation temperature was lowered to room temperature instead of 37 °C. With this new protocol, we still observed significant differences between the **SiR-glib**-treated samples and untreated controls (**Supporting Figure S1)** Notably, we verified our measurements at different developmental stages of the parasite in comparison to uninfected RBCs.

These results gave confidence to try **SiR-glib** in a potential diagnostic setup to be used in malaria endemic areas. As such, we turned to a battery-powered, portable smartphone-based microscope which uses a 180 mW 635 nm pen laser for excitation at a ∼45° angle, a smartphone as a camera and an 8-USD objective lens^[10]^. This setup is affordable, easy to assemble and was previously used for detection of single nucleic acid targets with the help of DNA origami nanoantennas^[10]^. The excitation wavelength of the device is similar to wavelengths of other malaria testing devices that have been used to detect the malaria pigment hemozoin^[24,25]^, and we wondered if **SiR-glib** may enhance the hemozoin-based detection of *P. falciparum*.

To get a first idea of the detection power of the smartphone microscope, we prepared uninfected RBCs and magnetically purified *P. falciparum* (FCR3 and 3D7) iRBCs with mature developmental stages, since these contain a larger amount of hemozoin. Both, RBCs and iRBCS were treated with 2 µM SiR-glib for 1 h. Afterwards, cells were washed and deposited between a glass slide and a glass coverslip to be investigated with the portable smartphone microscope versus untreated controls (**Figure 4A, B**). Videos were exported to single images of unbiased, different regions on which background subtraction was applied. Accordingly, the percentage of pixels above background threshold was counted. In vehicle-treated samples, we obtained images for uninfected, 3D7- and FCR3-iRBCs (**Figure 4C**), and after counting the pixels above threshold, we could not differentiate between non-infected and the 3D7 infected strain. Presumably due to higher auto-fluorescence stemming from the parasite, the FCR3-infected samples showed a significantly higher fluorescence, and could be clearly distinguished (**Figure 4D**). We performed the experiments in **SiR-glib** treated cells (**Figure 4E**) and found that 3D7- infected RBCs could now be significantly distinguished from non-infected cells (**Figure 4F**). FCR3 showed a comparable value to vehicle-treated controls, suggesting that the autofluorescence is either masking the **SiR-glib** signal, or that the staining protocol is less effective. In any case, the iRBCs showed significantly more pixels above the threshold and could thus be confirmed to carry the parasite.

**Figure 4.**
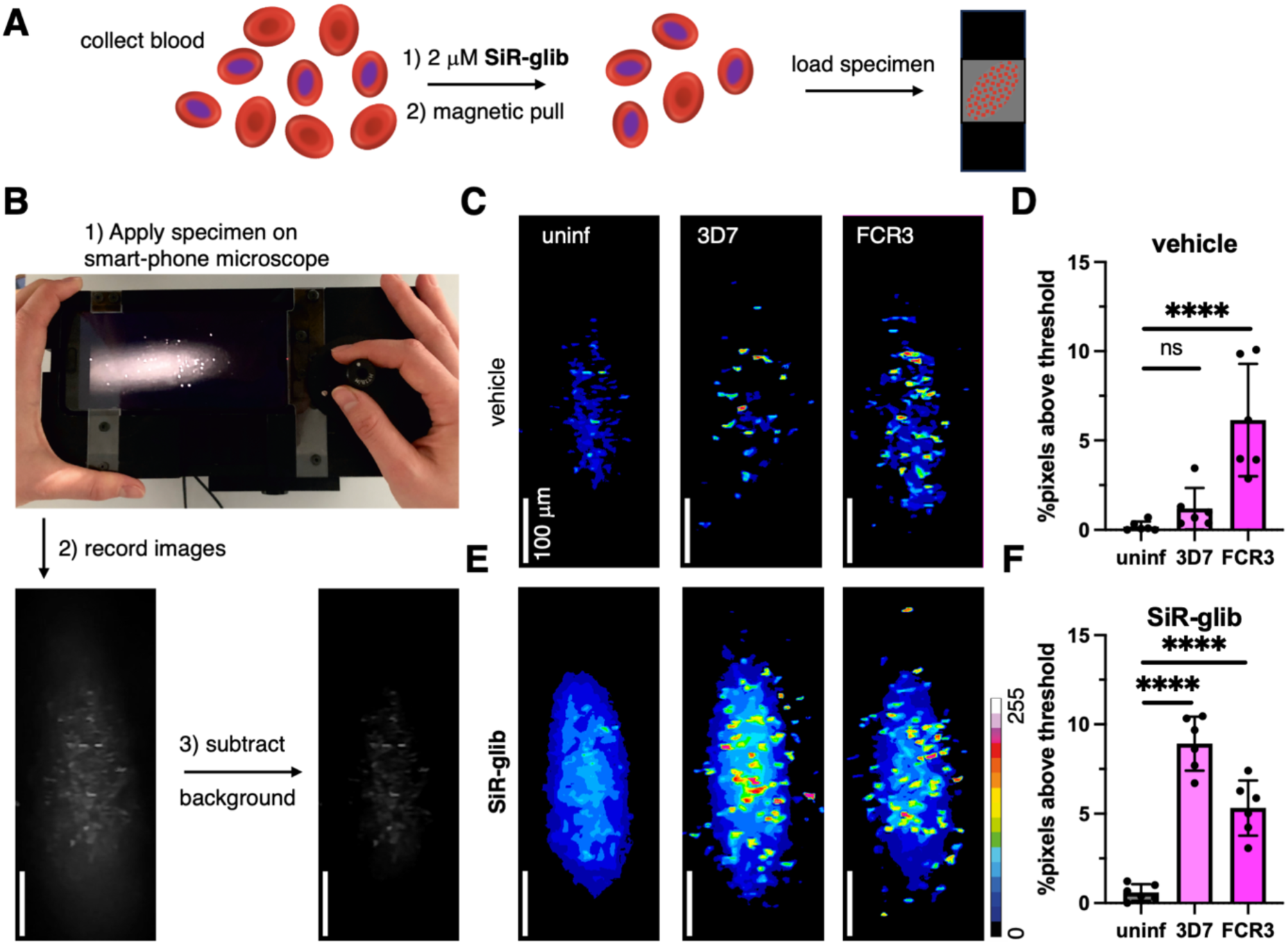
Imaging and analysis of *P. falciparum* infected RBCs using a portable smartphone-based microscope. **A)** Sample preparation for smart-phone based imaging. **B**) Workflow showing the mounted smartphone, image recording and background subtraction (photo reproduced from ref. ^[10]^). **C**) Vehicle controls of uninfected and 3D7- or FCR3-infected RBCs. **D**) Analysis of vehicle-treated RBCs show a significant increase in pixels above threshold for FCR3-infected RBCs. **E**) As for (**C**) but **SiR-glib**-treated RBCs. **F**) Analysis of **SiR-glib**-treated RBCs show a significant increase in pixels above threshold for RBCs infected with both strains, also enabling the detection of the 3D7 strain.

## Discussion

In this study, we demonstrate that **SiR-glib** specifically labels malaria-causing *P. falciparum* parasites in iRBCs, especially at the difficult to detect young ring stages. By optimizing the labelling protocol, we show that **SiR-glib** is able to identify parasite-containing RBCs within just 10 minutes. Lastly**, SiR-glib** is compatible with highly-portable smartphone**-**based microscopes, which can be deployed in areas without access to healthcare facilities and/or electricity, and with minimal training. As such, **SiR-glib** potentially provides a one-reagent point-of-care diagnostic that might be useful to identify and prioritize individuals with active malarial infection. Our successful approach pairing **SiR-glib** with an affordable and portable smartphone-based microscope creates a simple battery-powered, easy to apply diagnostic procedure that can be used at any location, with a first example of using this device on a cellular setting. The macro we provide for analysis can be run using FIJI (ImageJ), which is open source, without excessive training. While we acknowledge our diagnostic approach is currently not as fast as standard procedures used in malaria endemic areas^[1]^, this study marks a first step towards an alternative with high potential. For this reason, additional *P. falciparum* strains from endemic areas^[26]^ and other human-pathogenic *Plasmodium* species (such as *P. vivax*)^[5]^ need to be investigated and carefully compared.

Several diagnostic approaches utilize the hemozoin pigment to identify the parasite in patient blood^[4,15,27]^. However, the peripheral blood of malaria tropica patients exclusively contains ring stages and a few mature stage V gametocytes of *P. falciparum*. Only the latter developmental stage contains a high amount of hemozoin and is meant to be picked up by the *Anopheles* mosquito for sexual reproduction in the midgut. Ring stages are the most prominent developmental stage in the blood of a patient infected with *P. falciparum* and their hemozoin concentration is very low. Mature developmental stages of *P. falciparum* that contain a well-detectable amount of hemozoin, trophozoites and schizonts, sequester within the patients’ organs only appear within the peripheral blood after a splenectomy or when the patient is close to death. Interestingly, other human-pathogenic *Plasmodium* species do not sequester and all developmental stages of the parasite can be found in the peripheral blood of the patient. This important difference in parasite biology is used as one of the diagnostic criteria to determine which type of malaria the patient is suffering from.

The majority of diagnostic centers and rural diagnostic setups therefore still apply standard methods, which have been used for decades^[1]^. One of these methods is the Giemsa staining of thick or thin blood smears. Thick blood smears are usually used to determine the presence of a pathogenic organism while thin blood smears identify the (*Plasmodium*) species and the parasitemia (% parasitemia = (parasitized RBCs/total RBCs) × 100). Rapid Giemsa staining in a 10% solution is a procedure which takes up 10-15 min without taking the preparation of the blood film and the subsequent microscopic investigation by trained personnel into account^[1]^. Slow Giemsa staining using a 3% solution takes 45-60 min for the staining time only. For rapid diagnostic tests (RDTs), several kits are available for the detection of different antigens of *Plasmodium* species. These tests are mostly based on immunochromatography and are sold in cassette or dipstick format. Depending on the exact method, RDTs provide results in about 20 min^[28]^ and can be performed by trained personnel or the patient themselves. However, the recommended storage temperature for most RDTs is 4 °C, which requires refrigeration facilities that can limit take up in more rural (and tropical locations). The fact that **SiR-glib** can be stored and applied above room temperature for a long time and is easy to use are important requirements for the usage of the dye in a diagnostic field setting.

## Experimental Section

### Materials

Human blood and serum were obtained from the blood bank in Mannheim (Germany). If not otherwise indicated materials were obtained from: Gibco, c. c. pro GmbH, Thermo Fisher Scientific GmbH, Sigma-Aldrich, AppliChem, Carl-Roth GmbH & Co. KG, VWR Chemicals, Neofroxx GmbH and Serva. Chemicals were purchased from commercial vendors (Aldrich, TCI, Acros, etc.) and have been used without further purification.

### Synthesis of SiR-glib

Details on synthesis and chemical characterization are outlined in the Supporting Information.

### Mouse islet isolation and labeling

Male 8- to 12-week-old C57BL6 were socially housed in specific-pathogen free conditions under a 12 hour light-dark cycle, relative humidity 55 ± 10% and temperature 21 ± 2 °C. with ad libitum access to food and water. Animals were culled using a schedule-1 method and 1 mg/mL collagenase NB 8 (Serva) injected into the common bile duct, before digestion of the dissected pancreas in a water bath at 37 °C for 12 min with mild shaking. Islets were separated using gradient centrifugation in Histopaque-1119 and 1083 (Sigma-Aldrich). Islets were cultured in RPMI 1640 supplemented with 10% fetal bovine serum (FBS, Gibco), 100 units/mL penicillin, and 100 μg/mL streptomycin (Sigma-Aldrich), at 37 °C and 5% CO_2_.

Islets were incubated with 50 uM **SiR-glib** and 100 nM **LUXendin551** in complete medium for 1 hour at 37 °C and 5% CO_2_, before washing three times and imaging using a Zeiss LSM780 confocal microscope equipped with C-Apochromat 40x/1.20 W Korr M27 objective. Excitation and emission wavelengths for **SiR-glib** and **LUXendin551** were λex = 633 nm / λem = 639 – 692 nm and λex = 561 nm / λem = 571 – 649 nm, respectively. Animal studies were regulated by the Animals (Scientific Procedures) Act 1986 of the U.K. (Personal Project Licences P2ABC3A83 and PP1778740). Approval was granted by the University of Birmingham and University of Oxford Animal Welfare and Ethical Review Bodies (AWERB).

### Parasite strains

For this study the *P. falciparum* clones 3D7 and FCR3 were used. Both strains are laboratory model strains and regularly investigated in malaria research^[19,29,30]^. 3D7 is the limiting dilution clonal variant of the NF54 isolate^[20]^. NF54 was originally isolated from a malaria patient living near Amsterdam Airport Schiphol (so-called airport malaria) ^[21]^. The origin of the NF54 clone remains unknown. FCR3 was first isolated in Gambia and is a particularly dangerous strain for pregnant women^[31]^.

### Parasite culture

Parasites were kept in RPMI 1640 cell culture medium with 25 mM and HEPES *L*-glutamine (Gibco). The medium was supplemented with heat-inactivated type A+ human serum (5% (v/v)), AlbuMAX I (5% (v/v), Thermo Fisher Scientific GmbH), 20 µg/ml gentamycin (stock: 50 mg/ml, c. c. pro GmbH), 0.2 mM hypoxanthin (stock: 10 mM; c. c. pro GmbH). For our experiments the parasites were grown in culture with a hematocrit (HCT, volume percentage of RBC) of either 5% or 40%. A HCT of 5% is standard in cell culture, while a HCT of 40% is comparable to the HCT in the human body^[23]^. The *P. falciparum* 5% HCT cultures are maintained in 10 cm cell culture plates. 40% HCT cultures are kept in 6-well plates to avoid excess usage of RBC. The cultures are incubated in a cell culture incubation cabinet at 37 °C (CO_2_: 2.9%; O_2_: 5.8%; rH: 93%). The parasitemia of the *P. falciparum* cultures are assessed every day via thin methanol-fixed and Giemsa-stained blood smear^[4]^. The parasitemia of the culture was kept between 3-5%.

### SiR-glib labeling of infected RBCs from a 5% HCT culture

200 μL of a 5% hematocrit culture are required for **SiR-glib** labeling. The 200 μL are placed in a 1.5-ml reaction tube and centrifuged at 1,800 rpm for 30 seconds. The supernatant is discarded. Two washing steps each with 500 μL of cell culture medium and centrifugation for 30 seconds at 1,800 rpm are performed afterwards. The RBC pellet which remained after the second washing step is resuspended in 200 μL of cell culture medium supplemented with 2 μM **SiR-glib** or SiR (concentration of both stock solutions is 3.2 mM in DMSO). RBC which remained untreated were resuspended in 200 μL of cell culture medium. The cells are transferred to a 96-well microtiter plate, with one batch being divided into 2 wells so that each well contains approximately 100 μL. The cells are then incubated for 1 hour at 37 °C. Afterwards, RBC were washed twice with 500 μL of cell culture medium and centrifugation for 30 seconds at 1,800 rpm. The pelleted RBC are resuspended in 500 μL of cell culture medium and are now ready for microscopy. The sample is pipetted into a custom-made imaging chamber and examined^[14]^.

### SiR-glib labeling of infected RBCs from a 40% HCT culture

The same principle is used as for labeling a 40% HCT culture but with 400 μL of resuspended parasites from our cultures, placed in a 1.5-ml reaction tube. The washing steps are performed with 200 μL of cell culture medium and at the end of the procedure the pellet is suspended in 200 μL of cell culture medium for imaging.

### Short incubation of *P. falciparum*-infected RBCs with SiR-glib

For the short incubation with **SiR-glib**, two samples of 100 μL are taken from a 40% HCT culture. One sample is subsequently treated with **SiR-glib** and the other sample remains untreated as a negative control. The samples are centrifuged once at 1,800 rpm for 30 s. The supernatant is discarded and the pellet is resuspended in 100 μL RPMI cell culture medium supplemented with 50 µM **SiR-glib**. The pellet in the control sample is resuspended in 100 μL RPMI cell culture medium. The samples are incubated for 10 min at r.t. and immediately imaged.

### Image acquisition and analyses of *P. falciparum*-infected RBCs

Images (1024 x 1024 px) were acquired with the Axiovert 100 M/Zeiss CLSM 510 with a C-Apochromat 63x/1.2W corr objective. Samples were excited with the HeNe laser at a wavelength of 633 nm. The LP650 filter was used for the detection of the fluorescence. The fluorescence signal of parasitized and non-parasitized RBC is quantified with Fiji.

### Sample preparation for measurements on the smartphone microscope

A battery-powered smartphone microscope, previously described by Trofymchuk, Glembockyte et al. was used to demonstrate the detection with low-cost optical equipment^[8]^. Microscope cover slides (22 mm × 22 mm and 76 x 26 mm, Carl Roth GmbH, Germany) were cleaned using Ethanol 70%, dried with Kimtech Wipes (Merck KgaA, Germany) and 30 min treatment in UV-Ozone cleaner (PSD-UV4, Novascan Technologies, USA) at 100 °C. Dust was removed with compressed air. To create a flow chamber, two stripes of double-sided tape (3M, Germany) were glued onto the long edges of the large slide and the small cover slip was then laid on top. Mature parasites were purified using the MACS system (Miltenyi Biotec) as previously described^[29]^. 200 µl of iRBC were centrifuged for 30 seconds at 1,800 rpm at RT. The supernatant was discarded and the sediment was washed twice with 500 µl cell culture medium, centrifuging each washing step 30 s at 1,800 rpm RT. For staining, 200 µl of 2 µM **SiR-glib** in medium were added to the sediment and incubated for 1 h at 37 °C. Afterwards the sample was washed twice with 500 µl cell medium, performing centrifugation steps after each wash (30 s at 1,800 rpm, r.t.). For each sample, the sediment was diluted individually in medium to yield samples with similar blood cell concentration. The diluted blood sample was added to the chamber, which was sealed from one side with one Tough-Tag (Diversified Biotech) and closed with another from the other side.

### Measurements and analysis on the smartphone

Inside the home-built box a 638 nm laser diode with output power 180 mW (0638L-11A, Integrated Optics, UAB, Lithuania, driven by a portable power bank) was focused onto the sample at a ∼45° angle. After passing spectral filtering (BrightLine HC 731/137, Semrock Inc., USA), fluorescent signal was collected using an objective lens (NA = 0.25, LS-40166, UCTRONICS, USA) that guides the light to the monochrome camera of the smartphone (P20, Huawei, China). Movies were recorded via FreeDCam application (Troopii) and analyzed with ImageJ (Fiji). After file conversion with the FFMPEG plugin to .tif (32-bit), a home-written macro crops a defined region of interest in the video and calculates the area of pixels above a defined threshold. This threshold is individually set to the intensity value that is above the highest pixel intensities detected in the uninfected sample (100 for samples without dye, 120 with dye). The extracted data was analyzed using OriginPro2019, while the significance was determined using an ANOVA test.

### Statistical analyses

Statistical analyses of acquired data were conducted with SigmaPlot 13.0 and OriginPro2019.

### Image preparation and presentation

Microscopy images were prepared using Fiji.

## Conflict of interest

DJH and JB receives licensing revenue from Celtarys Research for provision of chemical probes. DJH has filed patents related to type 1 diabetes and type 2 diabetes therapy, unrelated to the present study.

## Acknowledgements

We thank Corentin Charbonnier, Jenny Eichhorst, and Ramona Birke (all FMP) for technical assistance. The authors would also like to thank Guillermo Gomez and Patrick Pitoniak for proofreading the manuscript. N. K. is grateful for Individual Funding from the Equal Opportunity Office at Heidelberg University. V.G. gratefully acknowledges financial support from the DFG (grant number GL 1079/1-1, project number 503042693). P.T. gratefully acknowledges financial support from the DFG (INST 86/1904-1 FUGG, excellence clusters NIM and e-conversion), BMBF (Grants POCEMON, 13N14336, and SIBOF, 03VP03891), as well as the Bavarian Ministry of Science and the Arts through the ONE MUNICH Project “Munich Multiscale Biofabrication”. This project has received funding from the European Union’s Horizon Europe Framework Programme (deuterON, grant agreement no. 101042046 to JB). DJH. was supported by MRC (MR/S025618/1), Diabetes UK (17/0005681 and 22/0006389) and UKRI ERC Frontier Research Guarantee (EP/X026833/1) Grants. This work was supported on behalf of the “Steve Morgan Foundation Type 1 Diabetes Grand Challenge” by Diabetes UK and SMF (grant number 23/0006627 to DJH and JB). This project has received funding from the European Research Council (ERC) under the European Union’s Horizon 2020 research and innovation programme (Starting Grant 715884 to DJH). This work was supported by grants from the European Research Council (ERC Advanced Grant 884281 “SynapseBuild” to VH) and the Deutsche Forschungsgemeinschaft (DFG, German Research Foundation; NeuroNex2/ HA2686/19-1 to VH). The research was funded by the National Institute for Health Research (NIHR) Oxford Biomedical Research Centre (BRC). The views expressed are those of the author(s) and not necessarily those of the NHS, the NIHR or the Department of Health. The project involves an element of animal work not funded by the NIHR but by another funder, as well as an element focused on patients and people appropriately funded by the NIHR.

## AUTHOR CONTRIBUTION

Conceptualization and Methodology: LR, KJ, VG, NK, JB; Formal analysis and investigation: CB, CC, IH, BG-F, ME, MW, SK, KR, CH, SG, JA, DR, ML, DJH, VG, NK and JB; Writing –Original Draft: DJH, NK, and JB; Reviewing and Editing: all authors; Visualization: VG, NK and JB; Supervision: VH, PT, KJ, VG, NK and JB; Funding Acquisition: DJH, PT, VG, and JB.

