## Supplemental Information for "A silicon rhodamine-fused glibenclamide to label and detect malaria-infected red blood cells"

**Affiliations**

**Table of Contents**


**1. General 4**

**2. Synthesis 5**

*2.1. 5-Chloro-2-methoxy-3-nitro-N-(4-sulfamoylphenethyl)benzamide (2) 5*

*2.2. 5-Chloro-N-(4-(N-(cyclohexylcarbamoyl)sulfamoyl)phenethyl)-2-methoxy-3-nitrobenzamide (3) 6*

*2.3. 3-Amino-5-chloro-N-(4-(N-(cyclohexylcarbamoyl)sulfamoyl)phenethyl)-2-methoxybenzamide (4) 7*

*2.4. 3-(6-Aminohexanamido)-5-chloro-N-(4-(N-(cyclohexylcarbamoyl)sulfamoyl)phenethyl)-2-methoxybenzamide (6) 8*

*2.5. 4-((6-((5-Chloro-3-((4-(N-(cyclohexylcarbamoyl)sulfamoyl)phenethyl)carbamoyl)-2-methoxyphenyl)amino)-6-oxohexyl)carbamoyl)-2-(7-(dimethylamino)-3-(dimethyliminio)-5,5-dimethyl-3,5-dihydrodibenzo[b,e]silin-10-yl)benzoate (SiR-glib) 9*

**3. NMR spectra 10**

*3.1. 5-Chloro-2-methoxy-3-nitro-N-(4-sulfamoylphenethyl)benzamide (2) 10*

*3.2. 5-Chloro-N-(4-(N-(cyclohexylcarbamoyl)sulfamoyl)phenethyl)-2-methoxy-3-nitrobenzamide (3) 11*

*3.3. 3-Amino-5-chloro-N-(4-(N-(cyclohexylcarbamoyl)sulfamoyl)phenethyl)-2-methoxybenzamide (4) 12*

*3.4. 3-(6-Aminohexanamido)-5-chloro-N-(4-(N-(cyclohexylcarbamoyl)sulfamoyl)phenethyl)-2-methoxybenzamide (6) 13*

*3.5. 4-((6-((5-Chloro-3-((4-(N-(cyclohexylcarbamoyl)sulfamoyl)phenethyl)carbamoyl)-2-methoxyphenyl)amino)-6-oxohexyl)carbamoyl)-2-(7-(dimethylamino)-3-(dimethyliminio)-5,5-dimethyl-3,5-dihydrodibenzo[b,e]silin-10-yl)benzoate (SiR-glib) 15*

### 1. General

All chemical reagents and anhydrous solvents for synthesis were purchased from commercial suppliers (Sigma-Aldrich, Fluka, Acros, Fluorochem, TCI) and were used without further purification if not stated otherwise. Commercial coumarin 461 and methylene blue were HPLC purified before concentration assessment and measuring photophysical properties to ensure similar purity and composition as their synthesizes, deuterated counterparts. BG-TMR and BG-SiR were described before.[1]

NMR spectra were recorded at 300 K in deuterated solvents on a Bruker AVANCE III HD 600 equipped with a CryoProbe or on Bruker AV-III spectrometers using either a cryogenically cooled 5 mm TCI-triple resonance probe equipped with one-axis self-shielded gradients or room temperature 5 mm broadband probe and calibrated to residual solvent peaks ($^{1}H/^{13}C$ in ppm): DMSO-$d_6$ (2.50/39.52), MeOD-$d_4$ (3.31/49.00). Multiplicities are abbreviated as follows: s = singlet, d = doublet, t = triplet, q = quartet, p = pentet, h = heptet, br = broad, m = multiplet. Coupling constants J are reported in Hz. Spectra are reported based on appearance, not on theoretical multiplicities derived from structural information.

LC-MS was performed on an Agilent 1260 Infinity II LC System equipped with Agilent SB-C18 column (1.8 µm, 2.1 × 50 mm). Buffer A: 0.1% FA in $H_2O$ Buffer B: 0.1% FA acetonitrile. The typical gradient was from 10% B for 0.5 min → gradient to 95% B over 5 min → 95% B for 0.5 min → gradient to 99% B over 1 min with 0.8 mL/min flow. Retention times ($t_R$) are given in minutes (min). Chromatograms were imported into Graphpad Prism8 and purity was determined by calculating AUC ratios.

Preparative or semi-preparative HPLC was performed on different instruments. An Agilent 1260 Infinity II LC System equipped with columns as followed: preparative column –Reprospher 100 C18 columns (10 µm: 50 x 30 mm at 20 mL/min flow rate; semi-preparative column – 5 µm: 250 x 10 mm at 4 mL/min flow rate. Eluents A (0.1% TFA in H2O) and B (0.1% TFA in MeCN) were applied as a linear gradient. Peak detection was performed at maximal absorbance wavelength. iii) A Waters e2695 system on a Supelco Ascentis® C18 HPLC Column (5 µm, 250 × 21.2 mm at 8 mL/min). Eluents A (0.1% TFA in $H_2O$) and B (0.1% TFA in MeCN) were applied as a linear gradient.

High resolution mass spectrometry was performed on an Agilent Technologies 6230 series accurate mass TOF LC-MS linked to an Agilent Technologies 1290 Infinity Series machine with a Thermo Accucore™ RP-MS column, 2.6 µm pore size, 30 × 2.1 mm, and a 3 min gradient from 5 to 99% aqueous MeCN with 0.1% TFA and MeCN with 0.1% TFA. flow rate: 0.8 mL/min; UV-detection: 220 nm, 254 nm, 300 nm.

### 2. Synthesis

#### 2.1. 5-Chloro-2-methoxy-3-nitro-*N*-(4-sulfamoylphenethyl)benzamide (2)

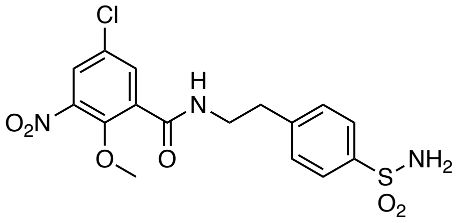


A round bottom flask was charged with 50 mg (0.216 mmol, 1.0 equiv.) 5-chloro-2-methoxy-3-nitrobenzoic acid (**1**) and dissolved in 1.2 mL DMSO and 97 µL (2.4 equiv.) DIPEA before 78.1 mg (0.25 mmol, 1.2 equiv.) of TSTU was added in one portion. The reaction mixture was stirred for 10 minutes before 51.8 mg (0.25 mmol, 1.2 equiv) of 4-(2-aminoethyl)benzenesulfonamide was added in one portion. The reaction mixture was stirred for additional 20 minutes before it was quenched with 10 mL of $dH_2O$, added dropwise under vigorous stirring. Finally, 1.2 mL of 1 M HCl was added to the reaction and the precipitate was collected by centrifugation at 4,000 rpm for 10 minutes. The supernatant was discarded, and the residue suspended in 10 mL $dH_2O$ / 1.2 mL 1 M HCl, and sedimentation was performed again. After removing the supernatant, the residue was transferred to a round bottom flask with MeOH and all volatiles evaporated to dryness. 72 mg (0.174 mmol) of the desired product was obtained in 81% yield as a yellowish powder.

**$^1$H NMR** (300 MHz, DMSO-$d_6$): $\delta$ [ppm] = 8.71 (t, $J$ = 5.5 Hz, 1H, NH), 8.16 (d, $J$ = 2.8 Hz, 1H), 7.75 (d, $J$ = 8.3 Hz, 2H), 7.73 (d, $J$ = 2.7 Hz, 2H), 7.46 (d, $J$ = 8.3 Hz, 2H), 7.32 (s, 2H, $NH_2$), 3.68 (s, 3H, $CH_3$), 3.51-3.59 (m, 2H), 2.92 (t, $J$ = 7.1 Hz, 2H).

**$^{13}$C NMR** (75 MHz, DMSO-$d_6$): $\delta$ [ppm] = 163.5, 148.4, 144.5, 143.4, 142.2, 134.0, 132.9, 129.2, 127.5, 125.7, 125.4, 63.2, 34.4. (one carbon signal is missing as it's overlapping with DMSO signal)

**HRMS** (ESI): calc. for $C_{16}H_{17}ClN_3O_6S$ $[M+H]^+$: 414.0521, found: 414.0511.

### 2.2. 5-Chloro-*N*-(4-(*N*-(cyclohexylcarbamoyl)sulfamoyl)phenethyl)-2-methoxy-3-nitrobenzamide (3)

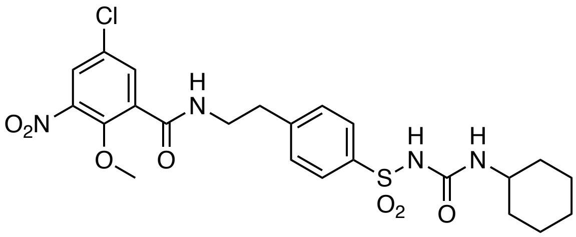


A round bottom flask was charged with 50 mg (121 µmol, 1.0 equiv.) of **2** and dissolved in 3 mL acetone before 33.2 mg (240 µmol, 2.0 equiv) of $K_2CO_3$ was added. The reaction mixture was heated to 70 °C and 23 µL (180 µmol, 22.6 mg, 1.5 equiv.) of cyclohexyl isocyanate diluted in 1 mL acetone was added dropwise. The reaction mixture was heated for an additional hour before 1 mL of 1 M HCl was added, and all volatiles were removed on a rotary evaporator. The residue was suspended in 10 mL d$H_2O$ / 1 mL 1 M HCl and the precipitate was collected after centrifugation at 4,000 rpm for 10 minutes. After removing the supernatant, the residue was transferred to a round bottom flask with MeOH and all volatiles evaporated to dryness. 64 mg (121 µmol) of the desired product was obtained in quantitative yield as a white powder.

**$^1$H NMR** (600 MHz, DMSO-$d_6$): $\delta$ [ppm] = 10.29 (s, 1H, NH), 8.67 (t, $J$ = 5.5 Hz, 1H, NH), 8.14 (d, $J$ = 2.7 Hz, 1H), 7.82 (d, $J$ = 8.3 Hz, 2H), 7.71 (d, $J$ = 2.7 Hz, 1H), 7.50 (d, $J$ = 8.3 Hz, 2H), 6.32 (d, $J$ = 7.9 Hz, 1H, NH), 3.61 (s, 3H, $CH_3$), 3.57 (q, $J$ = 6.5 Hz, 2H), 3.23-3.29 (m, 1H), 2.95 (t, $J$ = 7.0 Hz, 2H), 1.61-1.67 (m, 2H), 1.55-1.61 (m, 2H), 1.46-1.51 (m, 1H), 1.17-1.25 (m, 2H), 1.05-1.15 (m, 3H).

**$^{13}$C NMR** (150 MHz, DMSO-$d_6$): $\delta$ [ppm] = 163.4, 150.35, 148.3, 145.0, 144.45, 138.2, 133.9, 132.85, 129.25, 127.4, 127.25, 125.3, 63.0, 48.0, 34.3, 32.2 (2C), 24.9, 24.1 (2C).

**HRMS** (ESI): calc. for $C_{23}H_{28}ClN_4O_7S$ $[M+H]^+$: 539.1362, found: 539.1392.

### 2.3. 3-Amino-5-chloro-*N*-(4-(*N*-(cyclohexylcarbamoyl)sulfamoyl)phenethyl)-2-methoxybenzamide (4)

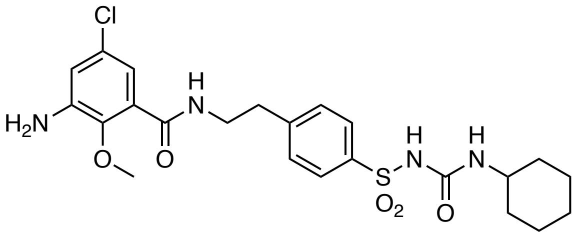

A round bottom flask was charged with 40 mg (74.3 µmol, 1.0 equiv.) of **3** under a nitrogen atmosphere and dissolved in 5 mL MeOH and 270 µL HOAc before 189 mg (12 mmol, 160 equiv.) of Zn dust was added in one portion. The reaction mixture was stirred for 1.5 hours at room temperature (solution turns red, and color fades over time) before it was centrifuged at 4,000 rpm for ten minutes to settle the remaining solids. The supernatant was collected and all volatiles were removed on a rotary evaporator. The crude was washed with 10 mL $dH_2O$, the solid filtered off and finally dried to obtain 34 mg (66.9 µmol) of the desired product in 90% yield as a white powder.

**$^1$H NMR** (600 MHz, MeOD-$d_4$): $\delta$ [ppm] = 7.91 (d, $J$ = 8.2 Hz, 2H), 7.51 (d, $J$ = 8.2 Hz, 2H), 6.86 (d, $J$ = 2.6 Hz, 1H), 6.84 (d, $J$ = 2.6 Hz, 1H), 3.69 (t, $J$ = 7.0 Hz, 2H), 3.49 (s, 3H, $CH_3$), 3.05 (t, $J$ = 7.0 Hz, 2H), 1.82-1.87 (m, 1H), 1.63-1.78 (m, 3H), 1.54-1.61 (m, 1H), 1.26-1.40 (m, 3H), 1.10-1.23 (m, 3H).

**$^{13}$C NMR** (150 MHz, MeOD-$d_4$): $\delta$ [ppm] = 168.0, 146.65, 144.7, 144.5, 140.1, 130.8, 130.6, 129.9, 128.8, 118.3, 117.8, 60.9, 50.1, 41.5, 35.9, 34.75, 33.8, 26.7, 26.5, 26.05, 25.8.

**HRMS** (ESI): calc. for $C_{23}H_{30}ClN_4O_5S$ $[M+H]^+$: 509.1620, found: 509.1625.

### 2.4. 3-(6-Aminohexanamido)-5-chloro-*N*-(4-(*N*-(cyclohexylcarbamoyl)sulfamoyl)phenethyl)-2-methoxybenzamide (6)

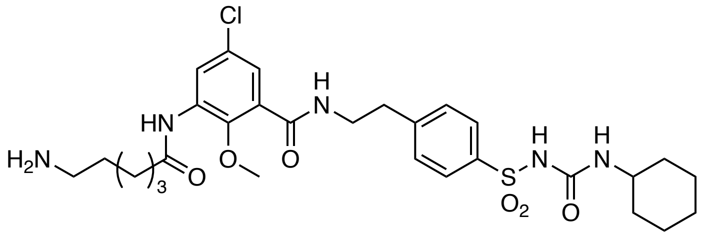


Under a nitrogen atmosphere, a round bottom flask was charged with 41 mg (117 µmol) of Fmoc-Ahx-OH and suspended in 1 mL $SOCl_2$. The reaction mixture was stirred for 1.5 hours at room temperature, and all volatiles were carefully removed under a stream of nitrogen in a well-ventilated fume hood. The residue was taken up in 1 mL of dry dioxanes, which was again carefully removed under a stream of nitrogen in a well-ventilated fume hood. Addition of 1 mL of dry dioxanes yielded a 117 mM stock solution, which was used immediately. A round bottom flask was charged with 10 mg (19.7 µmol, 1.0 equiv) of **4** dissolved in 2 mL of dry dioxanes and 13.3 µL of $NEt_3$, to which 505 µL of the acyl chloride (~3.0 equiv) was added dropwise under vigorous stirring. The reaction was stirred over night at room temperature before it was quenched with 1 mL of $dH_2O$ and all volatiles were removed in vacuo. The crude material was taken up in 800 µL DMF and 40 µL piperidine and allowed to incubate for 1 hour at room temperature, before it was quenched with 50 µL HOAc and 200 µL $dH_2O$ and subjected to RP-HPLC purification. The product containing fractions were pooled and 2.5 mg (3.40 µmol) of the desired product was obtained as a TFA salt after lyophilization in 17% yield over two steps as a white powder.

**$^1$H NMR** (600 MHz, DMSO-$d_6$): $\delta$ [ppm] = 10.34 (s, 1H, NH), 9.47 (s, 1H, NH), 8.43 (bs, 1H, NH), 8.16 (s, 1H), 7.83 (dd, $J$ = 8.2 Hz, $J$ = 1.8 Hz, 2H), 7.63 (bs, 2H, $NH_2$), 7.50 (dd, $J$ = 8.2 Hz, $J$ = 1.8 Hz, 2H), 7.12 (bs, 1H), 6.38 (d, $J$ = 6.8 Hz, 1H, NH), 3.55 (bs, 5H, $CH_2$ and $CH_3$), 3.23-3.31 (m, 1H), 2.95 (bt, $J$ = 6.8 Hz, 2H), 2.75-2.81 (m, 2H), 2.45 (bt, $J$ = 6.8 Hz, 2H), 1.51-1.67 (m, 8H), 1.44-1.51 (m, 1H), 1.30-1.37 (m, 2H), 1.16-1.26 (m, 2H), 1.07-1.15 (m, 3H).

**$^{19}$F NMR** (564 MHz, DMSO-$d_6$): $\delta$ [ppm] = -73.8.

**$^{13}$C NMR** (150 MHz, DMSO-$d_6$): $\delta$ [ppm] = 172.0, 164.6, 150.4, 146.4, 145.1, 138.2, 133.2, 131.2, 129.2, 127.2, 127.0, 123.0, 122.9, 61.4, 48.0, 40.0, 38.7, 35.7, 34.4, 32.2 (2C), 26.8, 25.4, 24.9, 24.5, 24.1 (2C).

**HRMS** (ESI): calc. for $C_{23}H_{27}ClN_4O_7S$ $[M+H]^+$: 622.2461, found: 622.2438.

#### 2.5. 4-((6-((5-Chloro-3-((4-(*N*-(cyclohexylcarbamoyl)sulfamoyl)phenethyl)carbamoyl)-2-methoxyphenyl)amino)-6-oxohexyl)carbamoyl)-2-(7-(dimethylamino)-3-(dimethyliminio)-5,5-dimethyl-3,5-dihydrodibenzo[*b*,*e*]silin-10-yl)benzoate (SiR-glib)

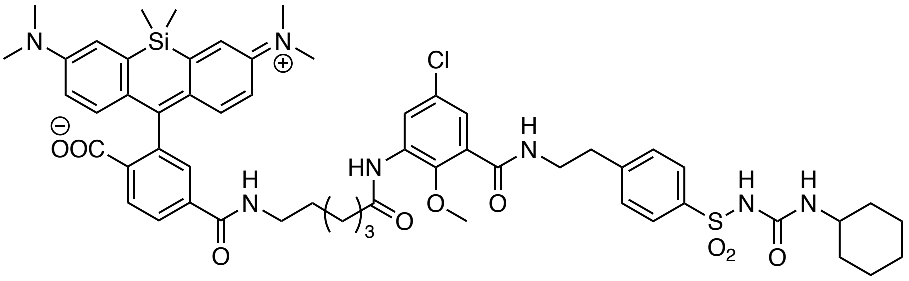

An Eppendorf tube was charged with 2.0 mg (4.24 µmol, 1.2 equiv.) of SiR-6-COOH and dissolved in 0.5 mL DMSO and 2.0 µL DIPEA before 1.6 mg (5.31 µmol, 1.5 equiv.) of TSTU was added in one portion. The reaction mixture was allowed to incubate for 10 minutes before it was added to an Eppendorf tube containing 2.6 mg (3.54 µmol, 1.0 equiv.) of **6** dissolved in 0.5 mL DMSO. The reaction mixture was vortexed and allowed to incubate for 2 hours, before it was quenched with 20 µL HOAc and 100 µL $dH_2O$ and subjected to RP-HPLC purification. The product containing fractions were pooled and 1.8 mg (1.50 µmol) of the desired product was obtained after lyophilization in 42% yield over two steps as a blue powder.

**$^1$H NMR** (600 MHz, DMSO-$d_6$): $\delta$ [ppm] = 10.29 (s, 1H, NH), 9.44 (s, 1H, NH), 8.43 (t, $J$ = 5.5 Hz, NH), 8.30 (d, $J$ = 8.5 Hz, 1H), 8.21 (d, $J$ = 8.5 Hz, 1H), 8.16 (bd, $J$ = 2.4 Hz, 1H), 7.88 (t, $J$ = 5.5 Hz, NH), 7.82 (d, $J$ = 8.3 Hz, 2H), 7.77 (bs, 1H), 7.50 (d, $J$ = 8.3 Hz, 2H), 7.12 (d, $J$ = 2.7 Hz, 1H), 7.04 (bs, 2H), 6.69 (bs, 4H), 6.32 (d, $J$ = 7.9 Hz, 1H, NH), 3.55 (m, 4H), 3.23-3.27 (m, 1H), 3.02-3.05 (m, 1H), 2.94 (bs, 12H), 2.87 (bs, 3H), 2.75 (bs, 1H), 2.43 (t, $J$ = 7.7 Hz, 1H), 2.29 (t, $J$ = 7.4 Hz, 1H), 1.95-2.02 (m, 1H), 1.62-1.67 (m, 1H), 1.56-1.59 (m, 2H), 1.40-1.49 (m, 2H), 1.19-1.32 (m, 4H), 1.08-1.14 (m, 1H), 0.94 (d, $J$ = 6.6 Hz, 1H), 0.85 (t, $J$ = 7.0 Hz, 1H), 0.62 (s, 3H), 0.53 (s, 3H).

**$^{19}$F NMR** (564 MHz, DMSO-$d_6$): $\delta$ [ppm] = -73.8.

**HRMS** (ESI): calc. for $C_{56}H_{66}ClN_7O_9SSi$ $[M+H]^+$: 1076.4173, found: 1076.4180.

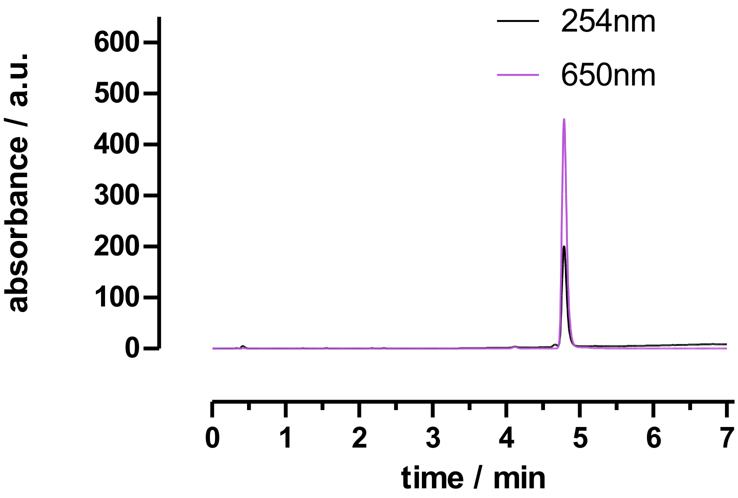

3. **NMR spectra**

**3.1. 5-Chloro-2-methoxy-3-nitro-*N*-(4-sulfamoylphenethyl)benzamide (2)**

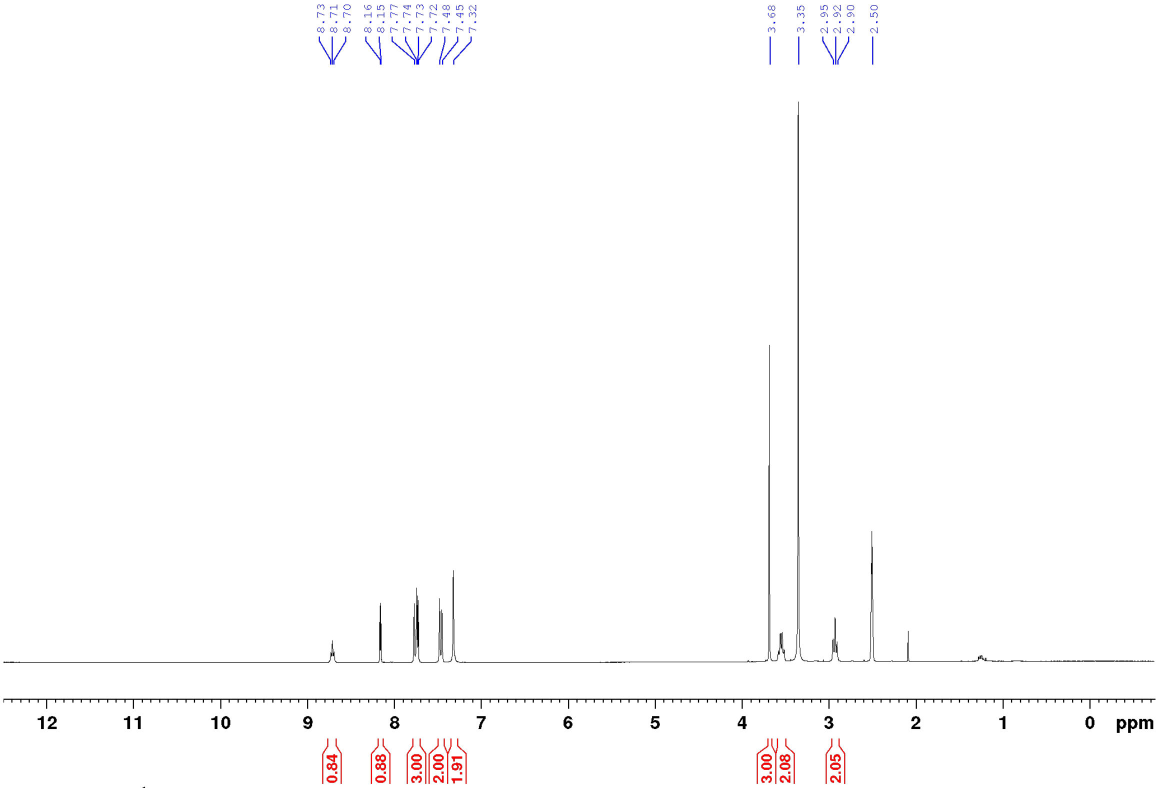


**Figure S1.** $^{1}H$ NMR spectra of compound **2** in DMSO-$d_6$.

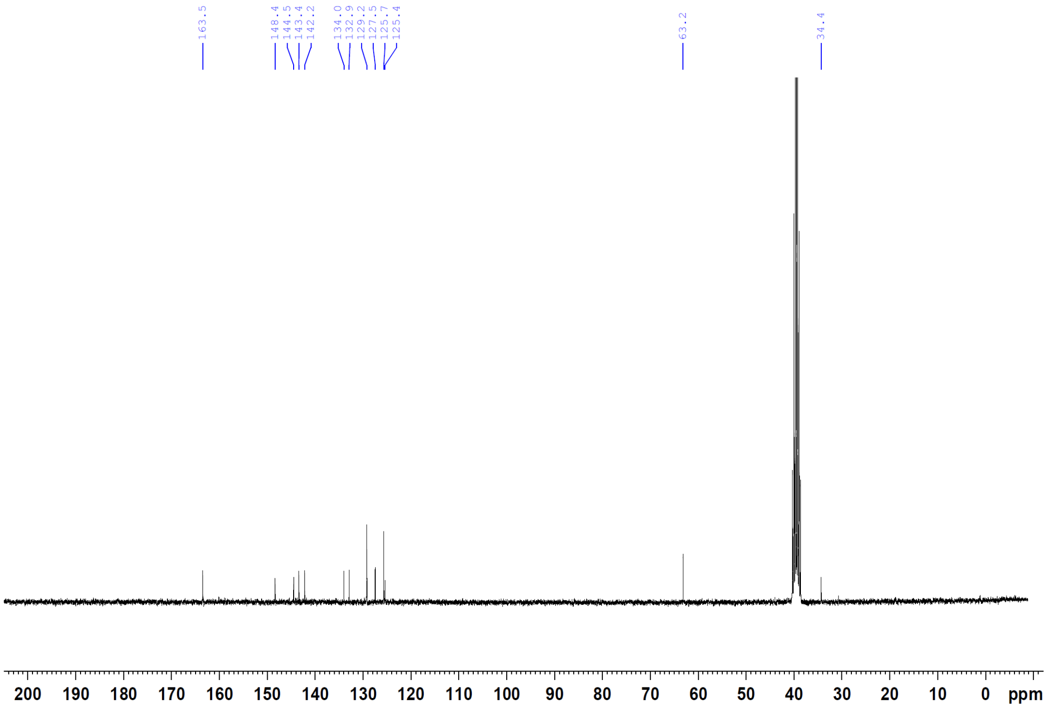


**Figure S2.** $^{13}C$ NMR spectra of compound **2** in DMSO-$d_6$.

### 3.2. 5-Chloro-*N*-(4-(*N*-(cyclohexylcarbamoyl)sulfamoyl)phenethyl)-2-methoxy-3-nitrobenzamide (3)

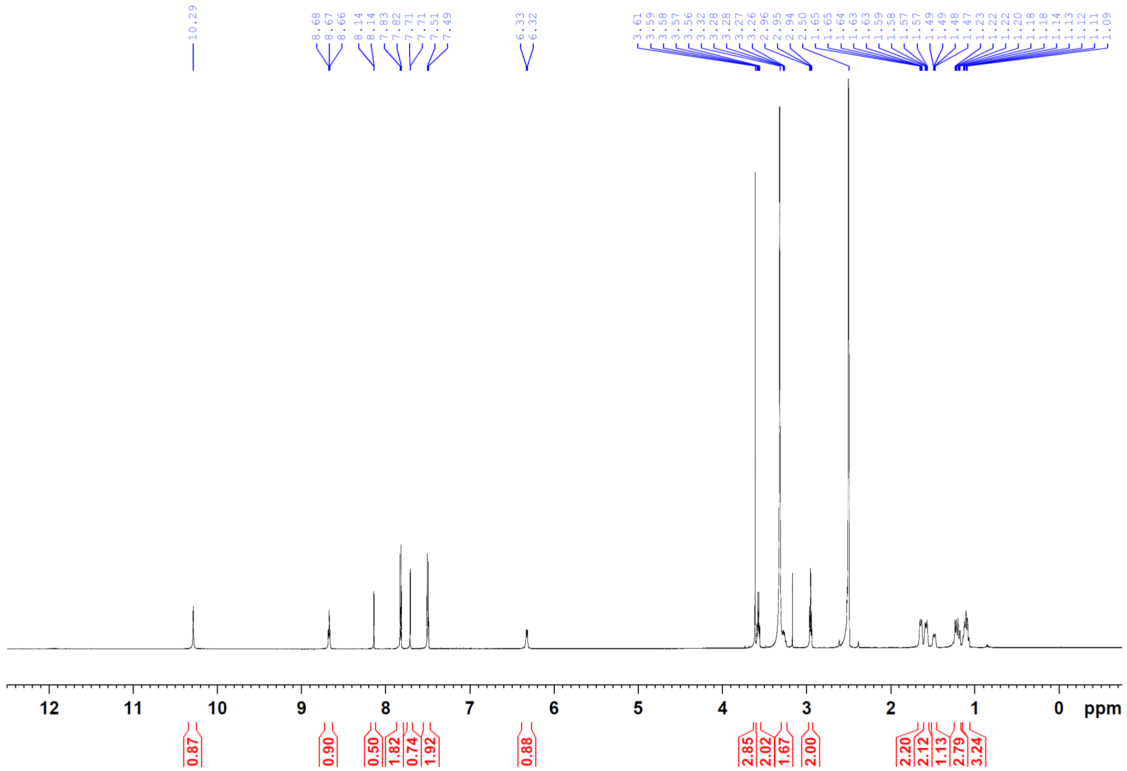


**Figure S3.** $^{1}$H NMR spectra of compound **3** in DMSO-$d_6$.

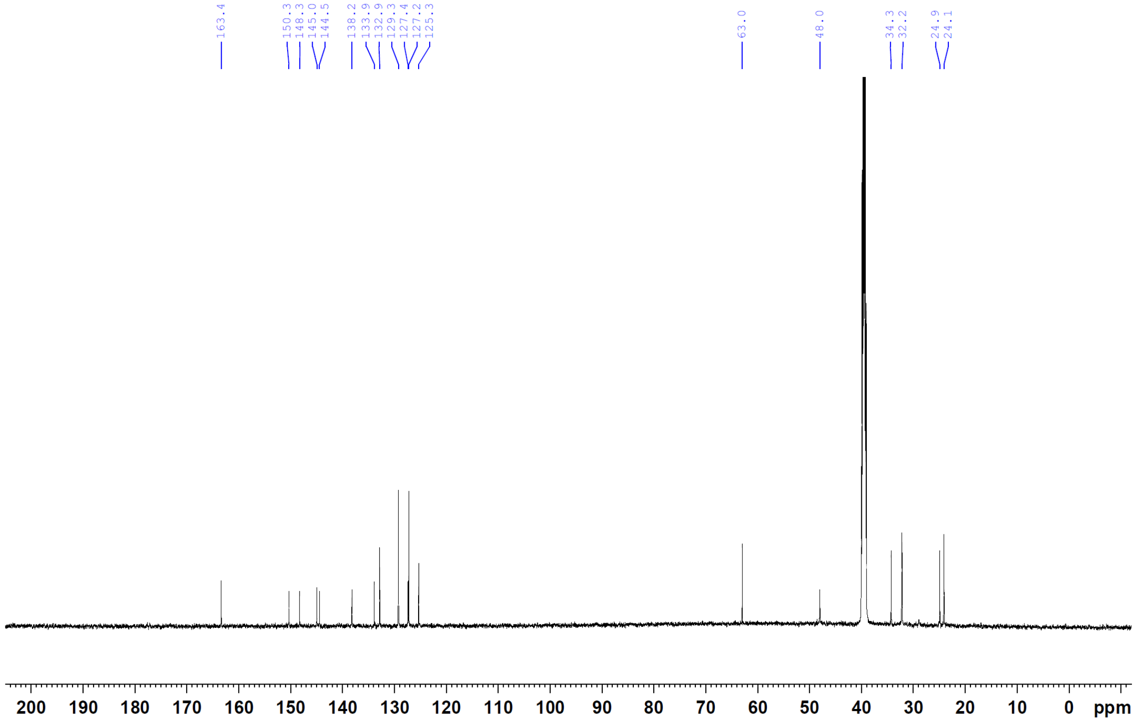


**Figure S4.** $^{13}$C NMR spectra of compound **3** in DMSO-$d_6$.

#### 3.3. 3-Amino-5-chloro-*N*-(4-(*N*-(cyclohexylcarbamoyl)sulfamoyl)phenethyl)-2-methoxybenzamide (4)

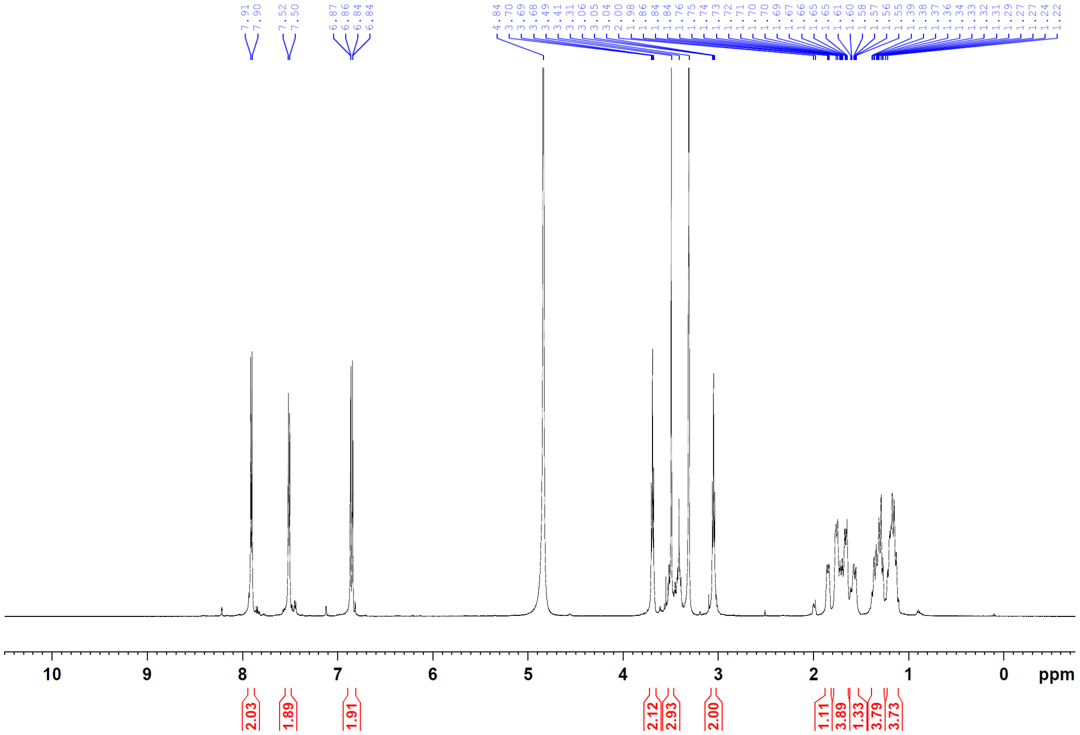


**Figure S5.** $^{1}H$ NMR spectra of compound **4** in MeOD-d4.

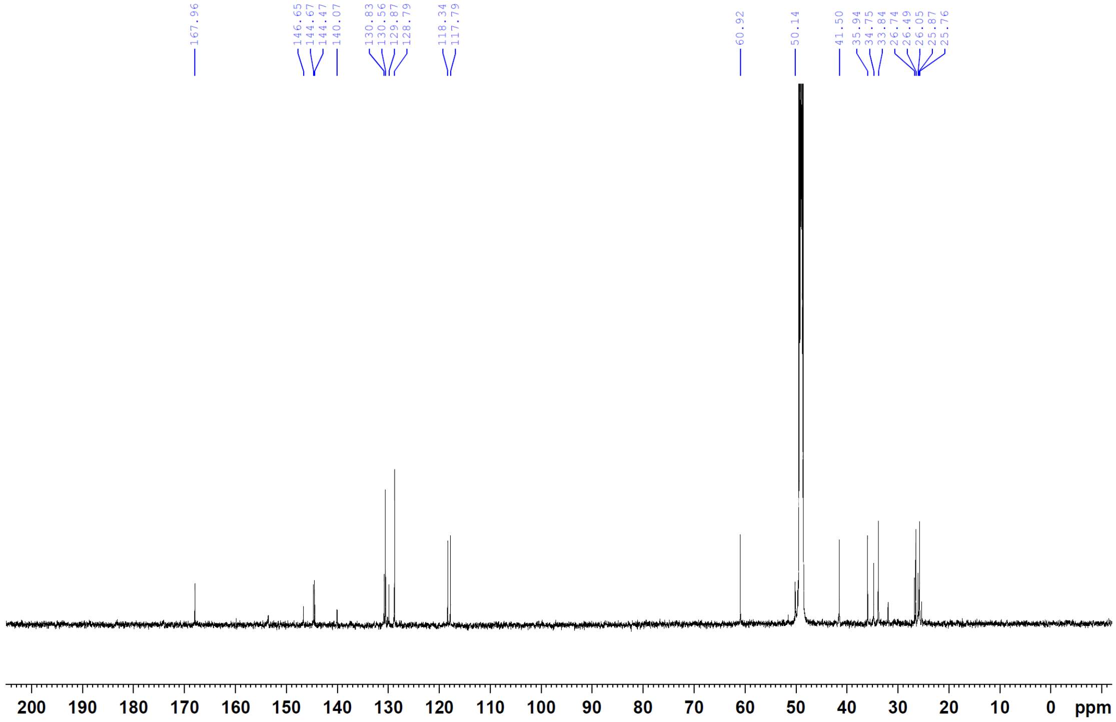


**Figure S6.** $^{13}C$ NMR spectra of compound **4** in MeOD-d4.

#### 3.4. 3-(6-Aminohexanamido)-5-chloro-*N*-(4-(*N*-(cyclohexylcarbamoyl)sulfamoyl)phenethyl)-2-methoxybenzamide (6)

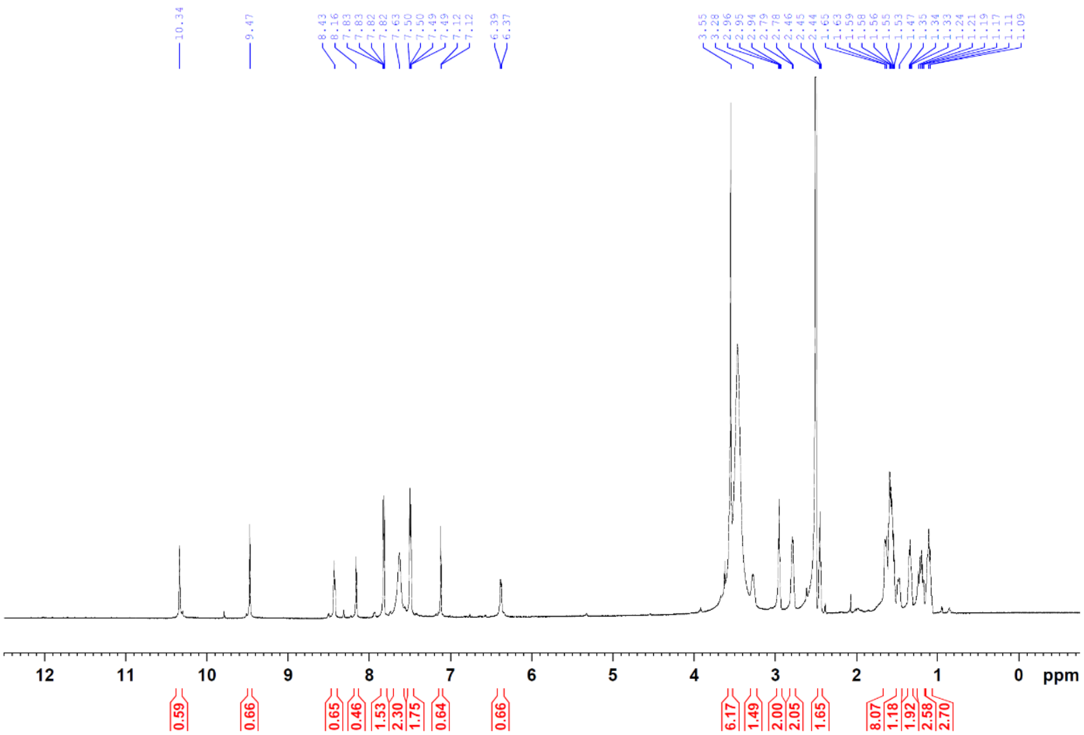


**Figure S7.** $^{1}$H NMR spectra of compound **6** in DMSO-$d_6$.

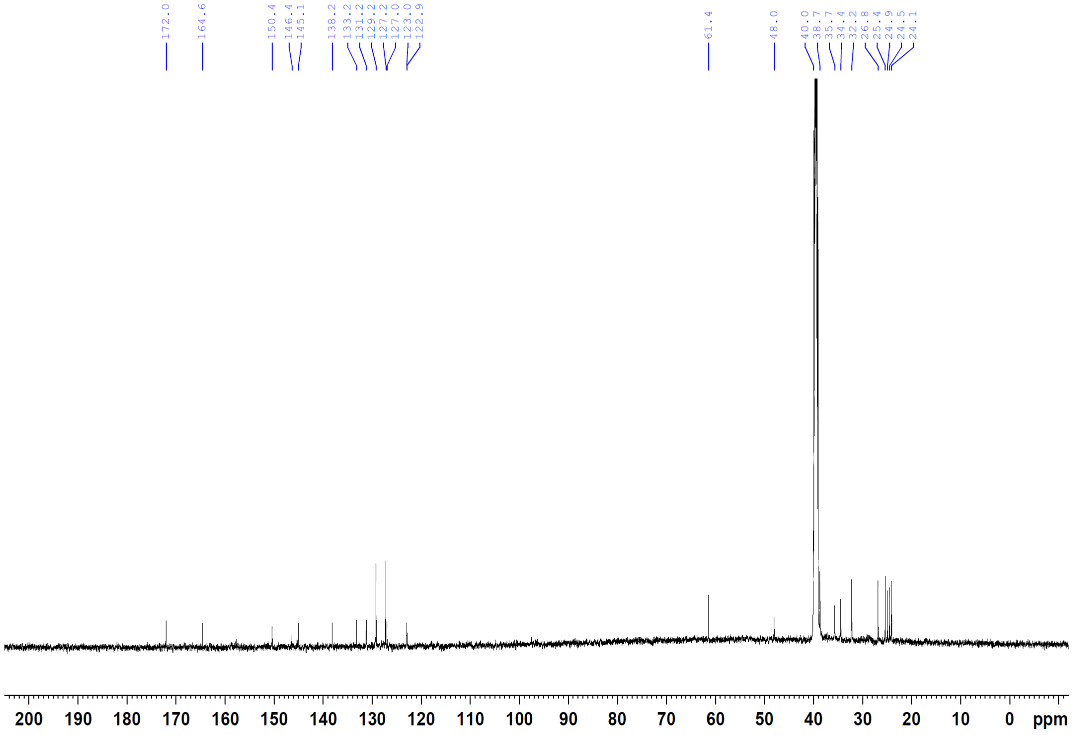


**Figure S8.** $^{13}$C NMR spectra of compound **6** in DMSO-$d_6$.

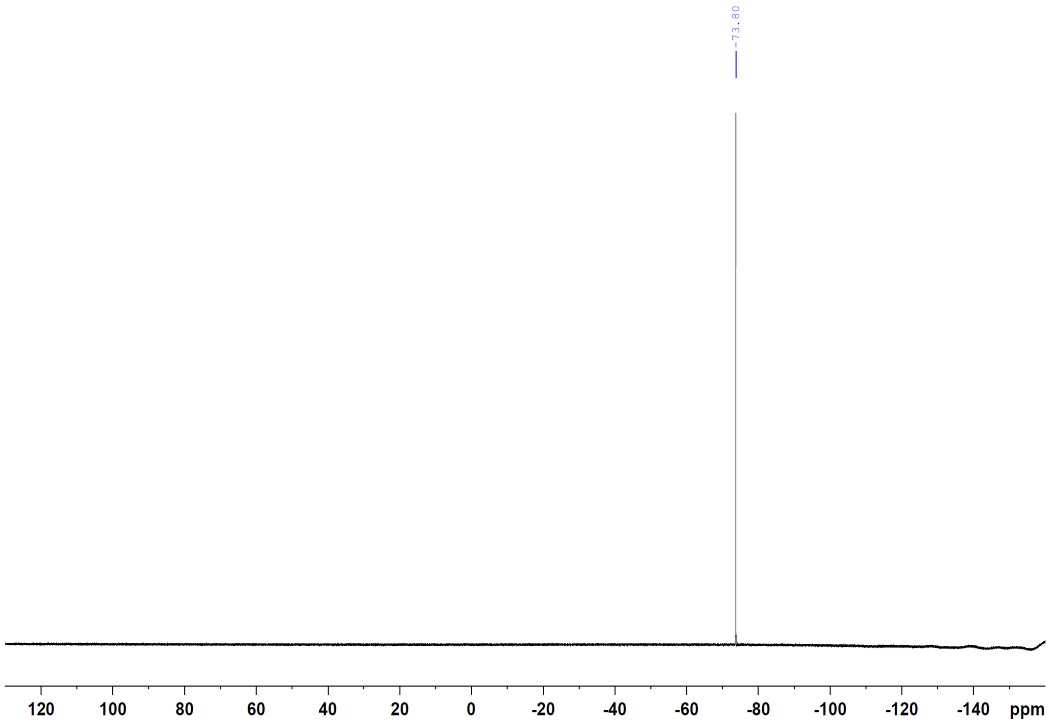


**Figure S9.** $^{19}F$ NMR spectra of compound **6** in DMSO-$d_6$.

#### 3.5. 4-((6-((5-Chloro-3-((4-(*N*-(cyclohexylcarbamoyl)sulfamoyl)phenethyl)carbamoyl)-2-methoxyphenyl)amino)-6-oxohexyl)carbamoyl)-2-(7-(dimethylamino)-3-(dimethyliminio)-5,5-dimethyl-3,5-dihydrodibenzo[*b*,*e*]silin-10-yl)benzoate (SiR-glib)

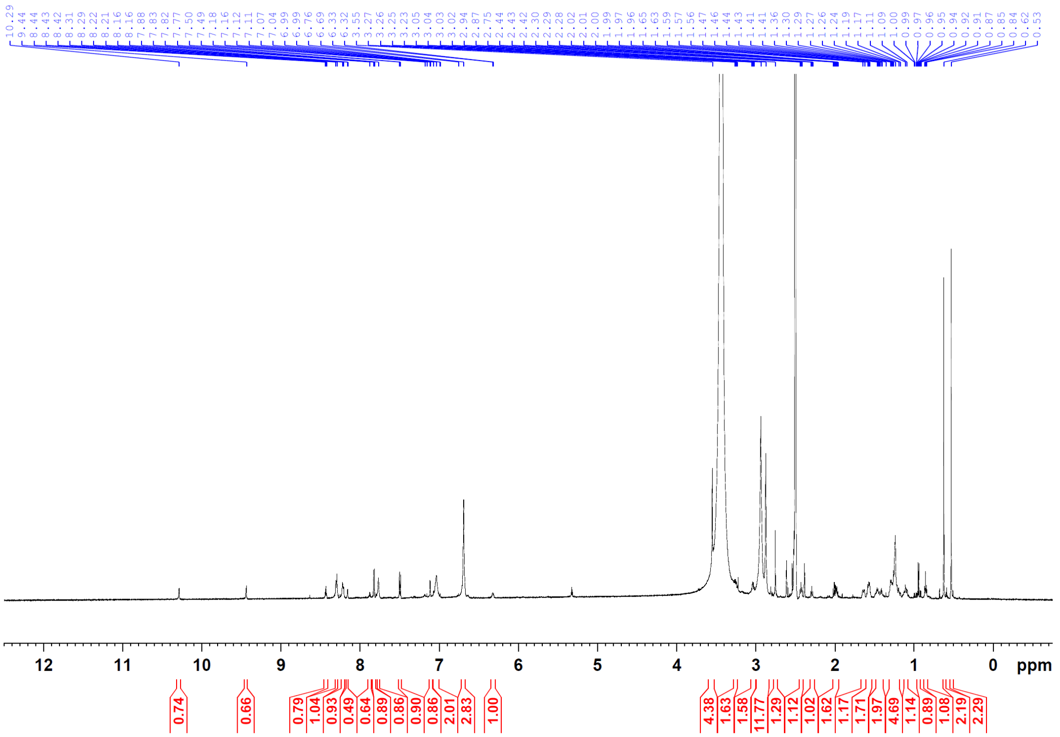


**Figure S10.** $^{1}H$ NMR spectra of compound **SiR-glib** in DMSO-$d_6$.

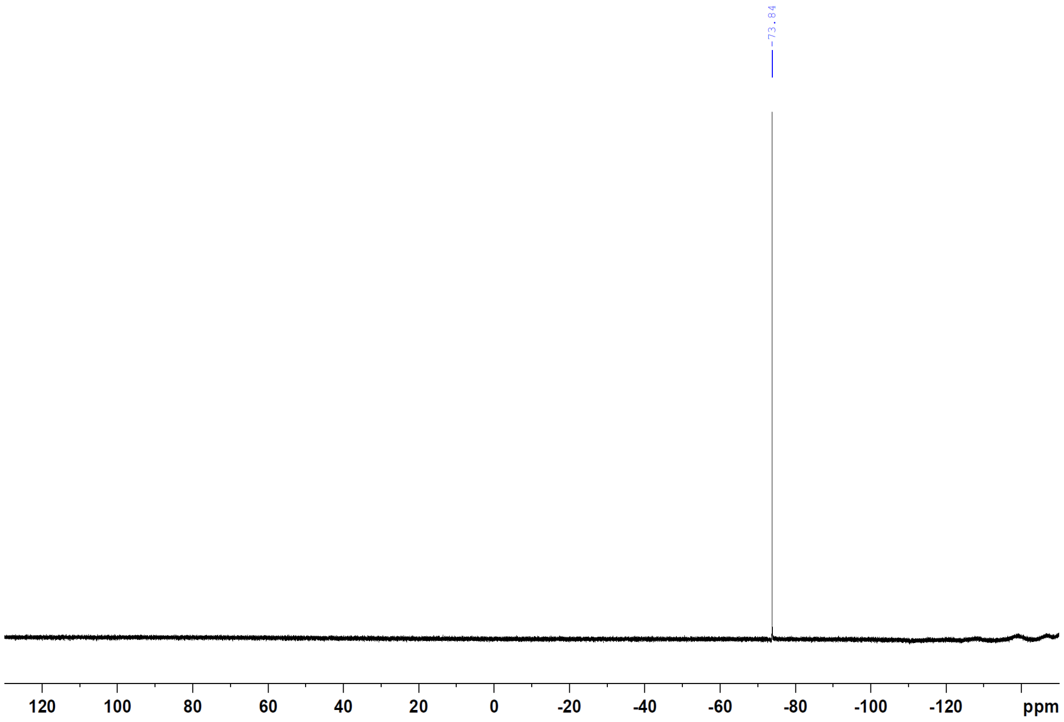


**Figure S11.** $^{19}F$ NMR spectra of compound **SiR-glib** in DMSO-$d_6$.
